# Distinct neural substrates of individual differences in components of reading comprehension in adults with or without dyslexia

**DOI:** 10.1101/2020.09.02.278267

**Authors:** O Ozernov-Palchik, TM Centanni, SD Beach, S May, T Hogan, JDE Gabrieli

## Abstract

Reading comprehension is a complex task that depends on multiple cognitive and linguistic processes. According to the updated Simple View of Reading framework, in adults, individual variation in reading comprehension can be largely explained by combined variance in three component abilities: (1) decoding accuracy, (2) fluency, and (3) language comprehension. Here we asked whether the neural correlates of the three components are different in adults with dyslexia as compared to typically-reading adults and whether the relative contribution of these correlates to reading comprehension is similar in the two groups. We employed a novel naturalistic fMRI reading task to identify the neural correlates of individual differences in the three components using whole-brain and literature-driven regions-of-interest approaches. Across all participants, as predicted by the simple view framework, we found distinct patterns of associations with linguistic and domain-general regions for the three components, and that the left-hemispheric neural correlates of language comprehension in the angular and posterior temporal gyri made the largest contributions to explaining out-of-scanner reading comprehension performance. These patterns differed between the two groups. In typical adult readers, better fluency was associated with greater activation of left occipitotemporal regions, better comprehension with lesser activation in prefrontal and posterior parietal regions, and there were no significant associations with decoding. In adults with dyslexia, better fluency was associated with greater activation of bilateral inferior parietal regions, better comprehension was associated with greater activation in some prefrontal clusters and lower in others, and better decoding skills were associated with lesser activation of bilateral prefrontal and posterior parietal regions. Extending the behavioral findings of skill-level differences in the relative contribution of the three components to reading comprehension, the relative contributions of the neural correlates to reading comprehension differed based on dyslexia status. These findings reveal some of the neural correlates of individual differences in the three components and the underlying mechanisms of reading comprehension deficits in adults with dyslexia.

## Introduction

Reading a connected text involves multiple cognitive and linguistic processes. According to the Simple View of Reading (Gough & Tunmer, 1986; Hoover & Gough, 1990), individual differences in reading comprehension are explained by two main skills: word decoding (i.e., accuracy of identifying words in print) and language comprehension (i.e., ability to understand spoken language). Although Decoding Accuracy and Fluency were not differentiated in the original model in children, the theory has since been expanded to include Fluency, measured by the rate of decoding words or naming stimuli, as an additional contributor, particularly in adulthood (Silverman et al., 2013; Tighe & Schatschneider, 2016; Tilstra et al., 2009). In the current manuscript, we differentiate between the two constructs and refer to accuracy as Decoding and fluency as Fluency. Support for the Simple View of Reading comes from numerous studies demonstrating that decoding and language comprehension are correlated but separable skills, together accounting for a large variance in reading comprehension performance across development (Aaron et al., 1999; Catts et al., 2003; de Jong & van der Leij, 2002; Hoover & Gough, 1990; Singer & Crouse, 1981). The current study examined the neural correlates of decoding, fluency, and language comprehension and examined whether those correlates were dissociable in the brain during naturalistic reading, as posited by the Simple View of Reading (SVR) framework.

An important prediction of the Simple View model is that the independence of word decoding and language comprehension will be particularly evident at the lower end of comprehension performance. Indeed, it has been shown that despite the positive and reciprocal association between decoding and language comprehension in typical readers (i.e., better decoding is associated with better language comprehension and vice versa), the correlation between the two skills weakens with lower decoding skills (Hoover & Gough, 1990; Singer & Crouse, 1981; Stanovich et al., 1984). Additionally, several studies have demonstrated that the relative contribution of language comprehension to reading comprehension increases with better decoding skills, both developmentally across grades (Catts et al., 1999, 2003, 2005; Francis et al., 2005; Hoover & Gough, 1990; Storch & Whitehurst, 2002; Vellutino et al., 1994) and across skill levels within one age group (Hoover & Gough, 1990).

An outstanding question exists, however, as to whether the relative contribution of the components to reading comprehension also differs in individuals with reading impairment. Developmental dyslexia (hereafter “dyslexia”) is an unexplained difficulty in learning to read, affecting approximately 10-12% of all individuals (Lyon, 1995). It is an outcome of multiple risk and protective factors (Ozernov-Palchik et al., 2016; Pennington & Ozonoff, 1996; van Bergen et al., 2014), but frequently originates from a deficit in phonological awareness (the ability to identify and manipulate speech sounds) (Stanovich & Siegel, 1994; Vellutino et al., 1996; Wagner et al., 1997). This difficulty results in slower and less accurate decoding. It has been suggested that slow fluency and labored decoding constrain comprehension during reading in individuals with dyslexia (Crain & Shankweiler, 1990), but since behavioral evidence is limited in differentiating the underlying mechanisms of performance, there has been insufficient evidence in support of this hypothesis. Given that the associations among the Simple View constructs differ based on decoding skill levels, it is plausible that the neurocognitive mechanisms underlying these skills, and the relations among them, will differ in individuals with dyslexia. For example, individuals with dyslexia have demonstrated greater reliance on language comprehension skills during reading, suggesting the use of linguistic context to bootstrap the decoding deficits that accompany this disorder (Nation & Snowling, 1998; Perfetti & Roth, 1980; Stanovich & Siegel, 1994). A compensatory account of dyslexia has been proposed that suggests increased reliance on language skills for reading comprehension in individuals with dyslexia to compensate for decoding difficulty (Snowling, 2005; Stanovich, 1980). Accordingly, reading comprehension will likely suffer in individuals with dyslexia who do not have these cognitive and linguistic resources available to them. Since similar reading comprehension performance and similar patterns of association among variables may have vastly distinct underlying neurocognitive mechanisms, using neuroimaging measures to study the SVR components separately in adults with and without dyslexia can reveal these differences.

Although much is known about the neurocognitive basis of impaired single-word decoding in dyslexia, very little is known about the neurocognitive basis of impaired reading comprehension in dyslexia. This represents a major gap in knowledge because it is the comprehension impairment that places the greatest burden on people with dyslexia throughout their education. The current study evaluated whether the proportional contributions of decoding, fluency, and language comprehension to reading comprehension performance, as well as the neural correlates of these components, differ in adults with dyslexia as compared to typical readers. The neural correlates of variation in the three skills have yet to be directly compared within the same individuals, but there are separate literatures examining which neural systems are related to decoding, fluency, and language comprehension performance. Better decoding skills have been associated with increased activation in the left temporoparietal, occipitotemporal, and inferior frontal regions (Martin et al., 2015), and reduced activation in these regions has been consistently demonstrated in individuals with dyslexia (Pugh et al., 2008; Richlan et al., 2011, 2013).

Language comprehension is supported by a distributed network of cortical regions including areas along the middle and superior temporal gyri and inferior frontal cortex that are known to play a role in language processing (Binder et al., 2009; Chai et al., 2016; Ferstl & von Cramon, 2001; Horowitz-Kraus et al., 2013; Huettner, 1989; Price, 2012), in addition to parietal, prefrontal, and posterior medial regions (e.g., cingulate cortex and precuneus) associated with attention and updating (Gernsbacher & Kaschak, 2003; Price, 2012; L. C. Robertson & Ivry, 2000; Roe et al., 2018; St George et al., 1999; Yarkoni, Speer, & Zacks, 2008b). Increased reading fluency has been associated with increased recruitment of the left-hemispheric occipitotemporal (i.e., visual word form) region (Benjamin & Gaab, 2012; Langer et al., 2015). To dissociate the neural substrates of individual variation in each skill, we directly compared the neural correlates of these skills within the same individuals.

To evoke reading comprehension processes, we designed a novel paragraph-reading task during imaging, which is a departure from the many prior studies employing single words or, more rarely, single sentences in isolation (Aboud et al., 2016; Richlan et al., 2011; Roe et al., 2018; Ryherd et al., 2018). Most of these tasks involved meta-cognitive judgments about words or sentences, such as making judgments about rhyming, spelling, or semantic plausibility (e.g., Price et al., 1997; Rimrodt et al., 2009). Despite the substantial contribution of these studies to understanding how the neural mechanisms that support reading differ in relation to specific skills, tasks requiring overt decisions or responses raise the likelihood that the neural responses observed reflect secondary, task-specific executive and motor demands in addition to reading processes. Such tasks, therefore, may be of limited utility in elucidating how the semantic, syntactic, and phonological processes interact in a naturalistic context to support reading comprehension (Rayner, 1998; Wehbe et al., 2014; Yarkoni, Speer, Balota, et al., 2008). Additionally, reading text sequentially, one word or a few words at a time, creates unnatural breaks and can hinder comprehension as it occurs during normal ecological conditions of reading connected text. Thus, a naturalistic paragraph reading design is ideal for investigating regions involved during comprehension in relation to individual differences in reading comprehension skills.

The current study applied a self-paced naturalistic fMRI task of reading text in 37 adults with and without dyslexia to examine the following: 1) whether decoding, fluency, and language comprehension make similar contributions to variance in reading comprehension in adults with or without dyslexia; 2) whether individual differences in decoding, fluency, and language comprehension are associated with distinct neural patterns of correlation during reading; 3) whether these neural correlates differ based on dyslexia status; 4) which of the neural correlates contributes to unique variance in reading comprehension skills across the groups and within each group.

This study is the first to investigate in-scanner connected-text reading in relation to individual differences in decoding, fluency, and comprehension skills. Using an unconstrained reading-aloud paradigm provides ecological validity as well as the ability to probe multiple aspects of reading as they unfold implicitly in natural reading. Based on the behavioral evidence, we hypothesized that decoding and fluency would be associated with distinct neural substrates of reading compared with language comprehension. We also predicted that decoding and fluency would involve the ventral reading systems and comprehension would involve brain regions implicated in semantic knowledge and cognitive control. Because of evidence of increased reliance on linguistic and cognitive systems in dyslexia, we expect that the neural correlates of the individual simple-view components would be different in individuals with dyslexia. Finally, we predicted that the neural substrates of each of the components would contribute to unique variance in predicting individual differences in reading comprehension.

## Methods

### Participants

Adults (n = 18 with dyslexia, n = 19 typical readers, age 18-41 years, M = 26.6, SD = 6.3) participated in the study. The original sample included 60 participants, but 12 participants were determined ineligible during the behavioral session, 3 participants were subsequently excluded due to incidental findings in MRI, and 8 participants (5 with dyslexia, 3 typical readers) were excluded due to an unacceptable number of scans with motion outliers (see Neuroimaging Processing). The demographic information for all participants is reported in Supplemental Table 1. All participants met eligibility criteria including: being a native speaker of American English; right-handed; born after at least 36 weeks gestation; no sensory or perceptual difficulties other than corrected vision; no history of head or brain injury or trauma; no neurological, neuropsychological, or developmental disorder diagnoses; no medications affecting the nervous system; nonverbal IQ standard score > 85 (Wechsler Adult Intelligence Scale Matrices subtest, Wechsler, 1981). Hearing tests were completed for all participants, and one participant with atypical hearing was excluded. The study was approved by the Committee on the Use of Humans as Experimental Subjects (COUHES) at the Massachusetts Institute of Technology. All participants provided informed written consent in order to participate.

### Behavioral Measures

All participants completed a comprehensive battery of standardized reading, language, and cognitive assessments, as well as a background questionnaire. In the current analyses, we included four standardized tests that capture the separate components of reading comprehension: decoding, fluency, and language comprehension. Reading comprehension comprised the outcome variable for the behavioral analyses. The decoding construct was measured by the Word Attack subtest of the Woodcock Reading Mastery Test - Revised/Normative Update (Woodcock, 2011); the language comprehension construct was measured by the Listening Comprehension subtest of WRMT-R/NU; and reading comprehension was measured by the reading comprehension subtest of the Gray Oral Reading Test-4 (GORT; Wiederholt et al., 2001). The fluency construct was measured by the Letters subtest of Rapid Automatized Naming (RANL; Wolf & Denckla, 2005) because letter naming does not involve semantic processing or word decoding. Additional standardized and background measures, including measures used to characterize participants as having dyslexia, are reported in Table 1. Participants were included in the dyslexia group (Dys) based on a report of life-long reading difficulty and/or clinical diagnosis and based on a performance below the 25th percentile on at least two out of four standardized subtests of timed or untimed word or nonword reading (Test of Word Reading Efficiency’s Sight Word Efficiency and Phonemic Decoding Efficiency (TOWRE; Torgesen et al., 2012); WRMT’s Word ID and Word Attack). Participants were included in the typical reader group (Typ) based on performance at or above the 25th percentile on all four of the above subtests.

**Table 1:** Participant characteristics for Typ and Dys groups. Bolded constructs indicate the measures used in the brain analyses. For all measures standard scores are reported.

| Construct | Typical | Dyslexia | Significance |
| --- | --- | --- | --- |
| Age (years) | $26.14 \pm 6.15$ | $27.07 \pm 6.53$ | n.s. |
| Nonverbal IQ | $114.28 \pm 13.41$ | $107.96 \pm 15.18$ | n.s. |
| Word ID | $108.75 \pm 7.55$ | $89.92 \pm 9.96$ | $< 0.001$ |
| **Word Attack** | $102 \pm 8.37$ | $77.25 \pm 9.61$ | $< 0.001$ |
| Sight Word Efficiency | $109.1 \pm 13.99$ | $89.15 \pm 9.38$ | $< 0.001$ |
| Phonemic Decoding | $104.33 \pm 7.62$ | $83.29 \pm 8.61$ | $< 0.001$ |
| Vocabulary | $114.44 \pm 7.77$ | $107.71 \pm 8.05$ | 0.003 |
| **Reading Comprehension** | $10.26 \pm 1.97$ | $7.77 \pm 1.48$ | $< 0.001$ |
| Blending Words | $12.17 \pm 2.62$ | $10.60 \pm 2.96$ | 0.045 |
| Elision | $9.9 \pm 1.88$ | $8.12 \pm 2.51$ | 0.005 |
| Memory for Digits | $10.65 \pm 2.11$ | $9.76 \pm 3.05$ | n.s. |
| **Language Comprehension** | $107.42 \pm 7.24$ | $101.39 \pm 7.41$ | 0.052 |
| **RAN-Letters** | $109.52 \pm 5.44$ | $101.61 \pm 5.93$ | $< 0.001$ |
| RAN 2-Set | $114.21 \pm 8.91$ | $103 \pm 7.25$ | $< 0.001$ |

### Neuroimaging Acquisition

Imaging was performed using a Siemens 3T MAGNETOM Trio, a Tim System (Siemens Medical Solutions, Erlangen, Germany), and a commercial Siemens 32 channel head coil. Structural data were collected using MPRAGE with 2530 ms TR, FOV = 256, 1 × 1 × 1 mm resolution. Functional data were collected with 3 × 3 × 3.6 mm resolution, 2000 ms TR, 30 ms TE, 90° flip angle, 64 x 64 base resolution, and 32 slices approximately parallel to the AC/PC line with coverage of the entire cortex. Prior to each scan, four images were acquired and discarded to allow longitudinal magnetization to reach equilibrium. PACE, an online prospective motion correction algorithm (Thesen et al., 2000), was implemented to reduce the effect of motion artifacts on functional data.

### Naturalistic Reading Task

The task consisted of three conditions, with seven 16-second blocks for each condition. Participants read expository (i.e., texts written to convey factual information on a topic) paragraphs out loud in their normal reading voice and rate inside the MRI scanner. Words in each paragraph were developed and matched based on the age of acquisition, frequency, and imageability. For the control blocks, participants verbally indicated whether arrows on the screen were pointing up or down (i.e., by saying “up” or “down”) to control for motor artifacts related to speech production. Participants’ speech was recorded with an MRI-compatible microphone. Participants were also presented with a fixation cross and were asked to keep still and relax during the fixation condition.

**Figure 1:**
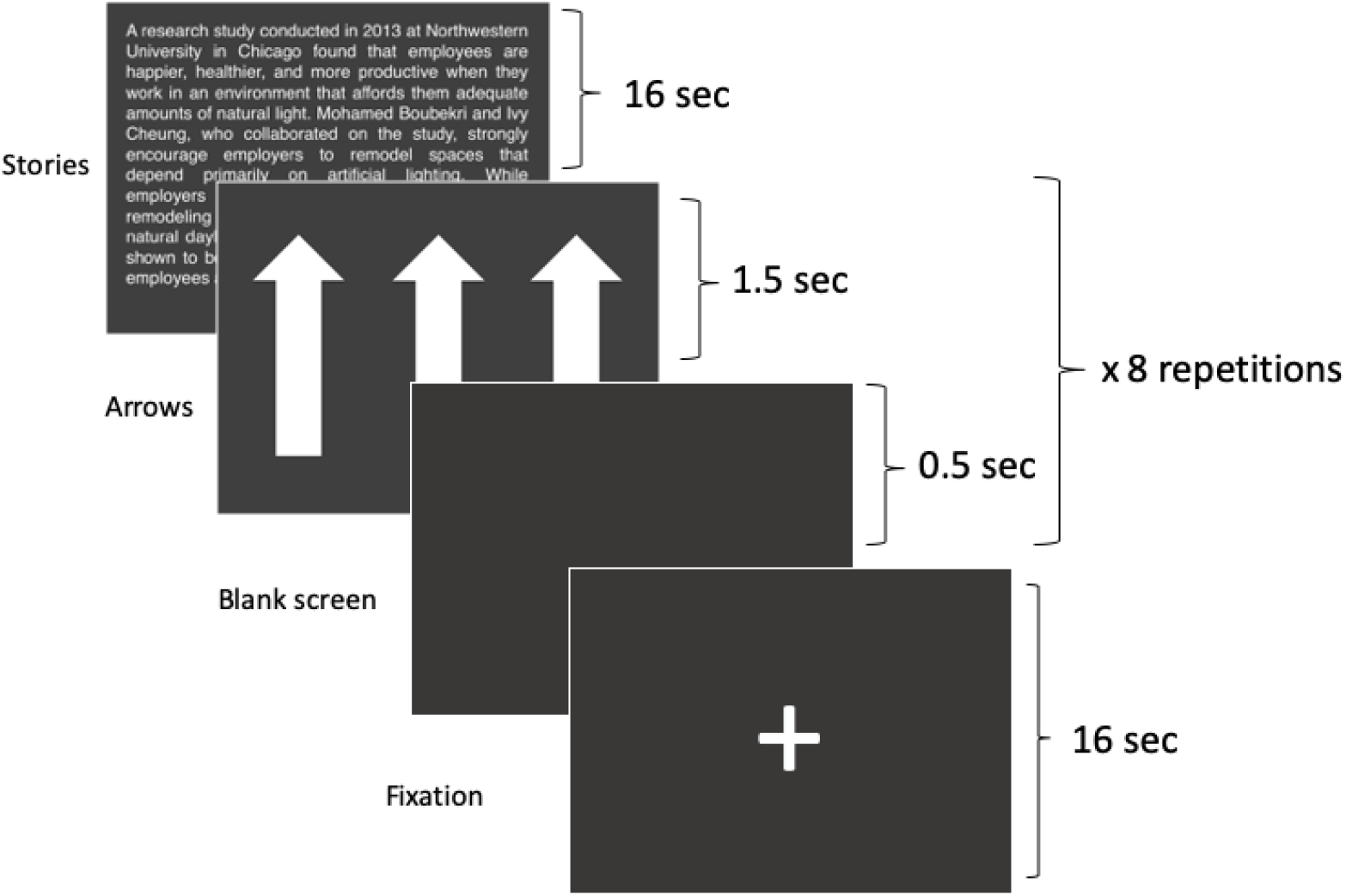
Timing of the scanner naturalistic reading task. The task included 7 blocks, with a randomized order of conditions. During the Arrow condition, the arrow screen alternating with a blank screen were repeated 8 times.

### Behavioral Analyses

We used multiple regression to examine whether the three SVR constructs (Language Comprehension, Word Attack, RAN Letters) contribute unique variance to reading comprehension. The models also included sex and age as control variables. The initial model included all participants who completed behavioral data (n = 60). The models in dyslexia and typical participants included only participants who were also included in the neuroimaging analysis (n = 37). The Shapiro test was used to evaluate whether variables violated the normality assumptions, and if the test was significant, permutated linear models were implemented using lmPerm (Wheeler et al., 2016). Multicollinearity among the variables in the regression models was evaluated using the variance inflation factor (VIF) from the olsrr package in R (Hebbali, 2018). In order to select the most parsimonious model in explaining reading comprehension from the three constructs, stepwise regression and variable relative importance analyses were conducted with 1,000 permutations.

### In-Scanner Performance

To ensure the validity of the in-scanner task, audio recordings of participants reading the stories inside the scanner were analyzed for all participants with adequate in-scanner recordings (n=36). Two measures were calculated: 1) task-fluency: a total score was derived from the total of number of words read for each of the stories; 2) task-accuracy: a mean score of number of words read accurately (using GORT scoring procedures) across the stories by a researcher blind to group assignment. To validate the in-scanner measure, pairwise Pearson correlation analyses were conducted relating the scores to the GORT accuracy and fluency measures.

### Neuroimaging Processing and Data Analysis

We preprocessed the fMRI data using FEAT (FMRI Expert Analysis Tool) Version 6.00, part of FSL version 5.0.2 (FMRIB’s Software Library, www.fmrib.ox.ac.uk/fsl) (Smith et al., 2004). High-resolution T1-weighted structural images were skull stripped. The functional data was then registered to the high-resolution structural image using Boundary Based Registration (BBR) algorithms (Greve & Fischl, 2009). We used FMRIB’s Linear Image Registration Tool (FLIRT; Jenkinson et al., 2002) to register the structural data to standard space (2mm MNI152). The following pre-modeling processing included the following steps: spatial smoothing with a Gaussian kernel of FWHM 5 mm; grand-mean intensity normalization by a single multiplicative factor; high pass temporal filtering. Statistical analysis was carried out with FILM using a double-gamma HRF model. The model included 6 motion regressors, their temporal derivatives, and nuisance regressors that modeled out single TR’s identified to have excessive motion according to a framewise displacement (FD) > 0.9mm (Siegel et al., 2014). Participants (*N* = 8) who lost more than 40% of frames (i.e., 70) due to FD censorship were excluded from the analysis. There were no significant group differences in the total number of outliers (*t*(31.7) = 1.42, *p* = 0.17) and the correlation between number of outliers and out-of-scanner GORT reading comprehension was not significant (*r*(34) = 0.07, *p* = 0.68). We carried out second-level analyses using a fixed effects model (Woolrich, 2008; Woolrich et al., 2004) with a cluster-forming threshold of z > 3.1 and a cluster probability of *p* < 0.05, using Gaussian random field theory (Worsley, 2001). Brain regions are reported in MNI coordinates and identified using the Harvard-Oxford atlas in the FMRIB software. For visualization of the statistical maps, we projected the data onto a standard volume using Mango software (http://rii.uthscsa.edu/mango). Coordinates and voxel size of group analysis results were obtained using the FSL version 5.0.2 Cluster tool.

### Whole-Brain Analyses

Across all participants, we ran a voxel-wise regression to examine the relation between Decoding (WA), Language Comprehension (LC), and Fluency (RANL), and whole-brain activation during passage reading versus arrows contrast. These scores were entered separately as covariates to a third-level group analysis using FLAME stage 1 (z > 3.1, P’s < 0.05; Woolrich, 2008; Woolrich et al., 2004) with Sex and Age as nuisance covariates. In order to identify the activation overlap at a group level among the three constructs, we performed a pair-wise conjunction analysis to determine the regions that were significantly active (z = 3.1) (Nichols et al., 2005).

### Whole-brain ROI Regression Analyses

We evaluated whether the regions associated with individual differences in each of the three constructs contribute uniquely and significantly to differences in reading comprehension. We extracted mean BOLD percent signal change for passages > arrows contrast from the clusters identified in the whole-brain regression models with each of the three constructs. The significant cluster values across participants were included in multivariate regression models using the same procedures as in the behavioral analyses.

### Literature-based Functional Regions of Interest (fROIs) Analyses-By Group

Based on the findings from the whole-brain analyses, regions of interest were selected from masks designed to delineate language and domain-general multi-demand systems (Table 2). The masks were downloaded from a publicly available website (https://evlab.mit.edu/funcloc). We used a set of 5 out of the 8 left-hemispheric language masks created using sentences > nonwords contrast (Fedorenko et al., 2010). We chose left-hemispheric masks corresponding to the four clusters identified in the whole-brain analyses for the three constructs: superior temporal (L_STG), angular gyrus (L_AngG), supramarginal gyrus (L_SMG), and lateral occipital lobe (L_LOcc). We also selected two prefrontal regions that, although they did not show a significant association in our whole-brain analysis, have been consistently reported in previous studies in relation to decoding and language comprehension skills: the opercular part of the inferior frontal gyrus (IFG) and inferior frontal gyrus orbitalis and its orbital part (L_IFGorb) (Martin et al., 2015; Price, 2012). Three masks from the right hemisphere homologues of the left-hemispheric language-selective regions were also used, including right angular gyrus (R_AngG), inferior frontal gyrus (R_IFG) and inferior frontal gyrus OB (R_IFGorb). It is still unclear whether the prefrontal regions recruited serve broader cognitive functions or whether they are part of the language network (Hancock et al., 2017). To delineate these possibilities, we included two prefrontal fROIs from the multiple demand (MD) system: posterior parietal (PostParietal) and the insula (Insula). The masks were derived using a hard > easy spatial working memory contrast (Blank et al., 2014). Finally, we selected an additional reading-specific mask from the Neurosynth database for the left putative visual word form area (VWFA). Mean activation values for the significant voxels for the stories > arrows contrast were extracted for each participant using the masks. We fit a permuted linear regression model using the lmPerm package (Wheeler & Torchiano, 2010) predicting each of the three skills from the literature-based fROIs across both Typ and Dys groups as follows:

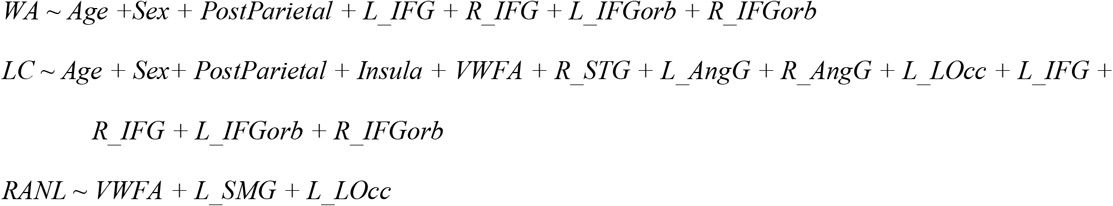

For each of the six models, we implemented a stepwise regression analysis to identify the optimal set of predictors for each construct separately in the Typ and the Dys groups. Predictors with high Variance Inflation Factors (VIF) were excluded from each model on a model-by-model basis.

## Results

### Behavioral

The two groups were similar in terms of sex (10 females in Typ and 11 females in Dys) and age (*p*’s > 0.98). There were significant differences in age between male and female participants (*t*(40.57) = −2.48, *p* = 0.02), with older male participants. Additionally, across both groups, males performed above females on many of the reading behavioral measures. Sex differences on measures of language and reading, as well as in the brain, have been reported across multiple studies (e.g., Lietz, 2006; Reilly, Neumann, & Andrews, 2019; Rutter et al., 2004). This topic is beyond the scope of the current study but is discussed elsewhere (e.g., Krafnick & Evans, 2017). We therefore included Age and Sex in the models.

Age, Sex, and the three constructs of Language Comprehension (LC), Fluency (RANL), and Decoding (WA), were combined in a stepwise regression model with 1,000 permutations to identify whether, in accordance with the Simple View of Reading, each of the three constructs contributed independent and significant variance in explaining reading comprehension. The R^2^ for the entire model was 0.49 (*p* < .001) and the best set of predictors included the three constructs in order of their relative importance: (1) Language Comprehension (contributing 41% of R^2^, ηp^2^ = 0.151), (2) Decoding (contributing 38% of R^2^, ηp^2^ = 0.139), and (3) Fluency (contributing 19% of R^2^, ηp^2^ = 0.177). Correlations among the three constructs revealed a significant positive association between Fluency and Decoding (*r*(52) = 0.6, *p* < 0.001), a nonsignificant association between Fluency and Language Comprehension (*r*(52) = 0.14, *p* = 0.3), and a borderline significant association between Language Comprehension and Decoding (*r*(52) = 0.26, *p* = 0.06). Although the three constructs were moderately correlated, the VIF values for all variables ranged from 1.09 to 1.64, suggesting that multicollinearity is not an issue of concern for these regression models (O’brien, 2007).

To test whether the constructs of reading comprehension are different based on dyslexia status, the models described above were repeated in the Dys and Typ groups in the final neuroimaging sample (n = 18 and 19, respectively, excluding participants who were not eligible to be part of either of the two groups based on their reading performance). For Typ, the overall model explained 36.3% of total variance in reading comprehension, with Language Comprehension accounting for 82% (ηp^2^ = 0.26) of total variance and making the only significant contribution to the model. For Dys, the model accounted for 28.3% of total variance with Language Comprehension contributing 28% (ηp^2^ = 0.189), Decoding, 8% (ηp^2^ = 0.04); and Fluency, 5% (ηp^2^ = 0.08) of total variance.

### In-Scanner Performance

Correlational analyses confirmed the construct validity of the in-scanner task. Task-fluency scores were significantly associated with GORT fluency scores (*r*(36) = 0.73, *p* < 0.001) and task-accuracy scores were significantly associated with GORT accuracy (*r*(36) = 0.65, *p* < 0.001). Additionally, both in-scanner measures significantly correlated with GORT comprehension scores (accuracy: *r* (36) = 0.52, *p* < 0.001; fluency: *r* (36) = 0.65, *p* < 0.001).

### Whole-Brain

To examine the activation induced by the reading task, we conducted a one-sample *t*-test for stories > arrows for the entire group (Figure 2A). This contrast yielded significant activation in the left-hemisphere ventral occipitotemporal and inferior frontal regions (occipital fusiform gyrus, inferior temporal gyrus, inferior frontal gyrus). Additional activations included right middle/superior frontal regions, basal ganglia (caudate, putamen), and paracingulate and cingulate regions. The Typ group, as compared to Dys group, showed significantly greater activation in left occipitotemporal region, which includes the putative VWFA (Figure 2B). There were no significant clusters with greater activation in Dys as compared to Typ.

**Figure 2:**
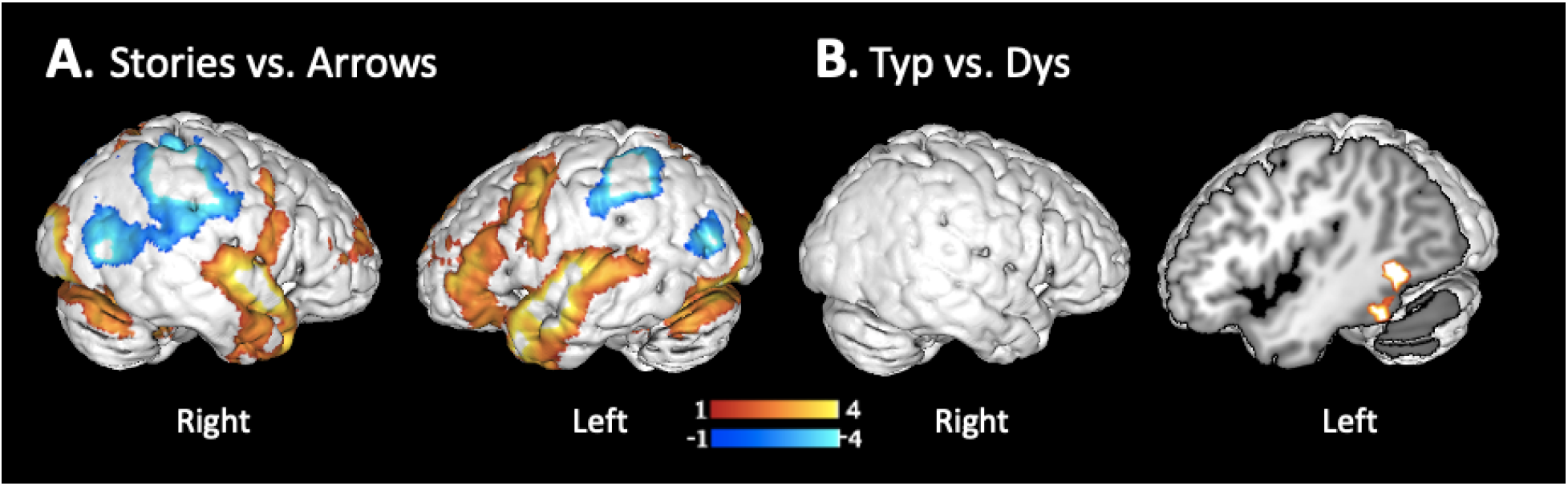
A) Whole brain reading task results for reading stories versus naming arrows contrast for all participants. Clusters for reading > arrows shown in yellow/red; for arrows > reading shown in blue. B) Group comparison for Typ versus Dys groups. For all results z-score thresholds represented by the colored bars. Z’s > 3.1, p’s < 0.05, corrected for multiple comparisons.

For the whole-brain correlation with each of the three constructs, there were significant negative correlations with Language Comprehension and Decoding, and a significant positive correlation with Fluency (Table 2, Figure 3). There were no significant clusters for the opposite relationships. Better Language Comprehension was significantly associated with less activation in bilateral posterior temporal and inferior parietal (including angular gyrus) and posterior parietal regions, insula, precuneus, and putamen. Better Decoding was significantly associated with less activation in medial and lateral posterior parietal cortex, including in the cuneus and precuneus. Finally, better Fluency was significantly associated with greater activation in left dorsal temporoparietal and ventral occipitotemporal regions, including the visual word form area.

**Figure 3:**
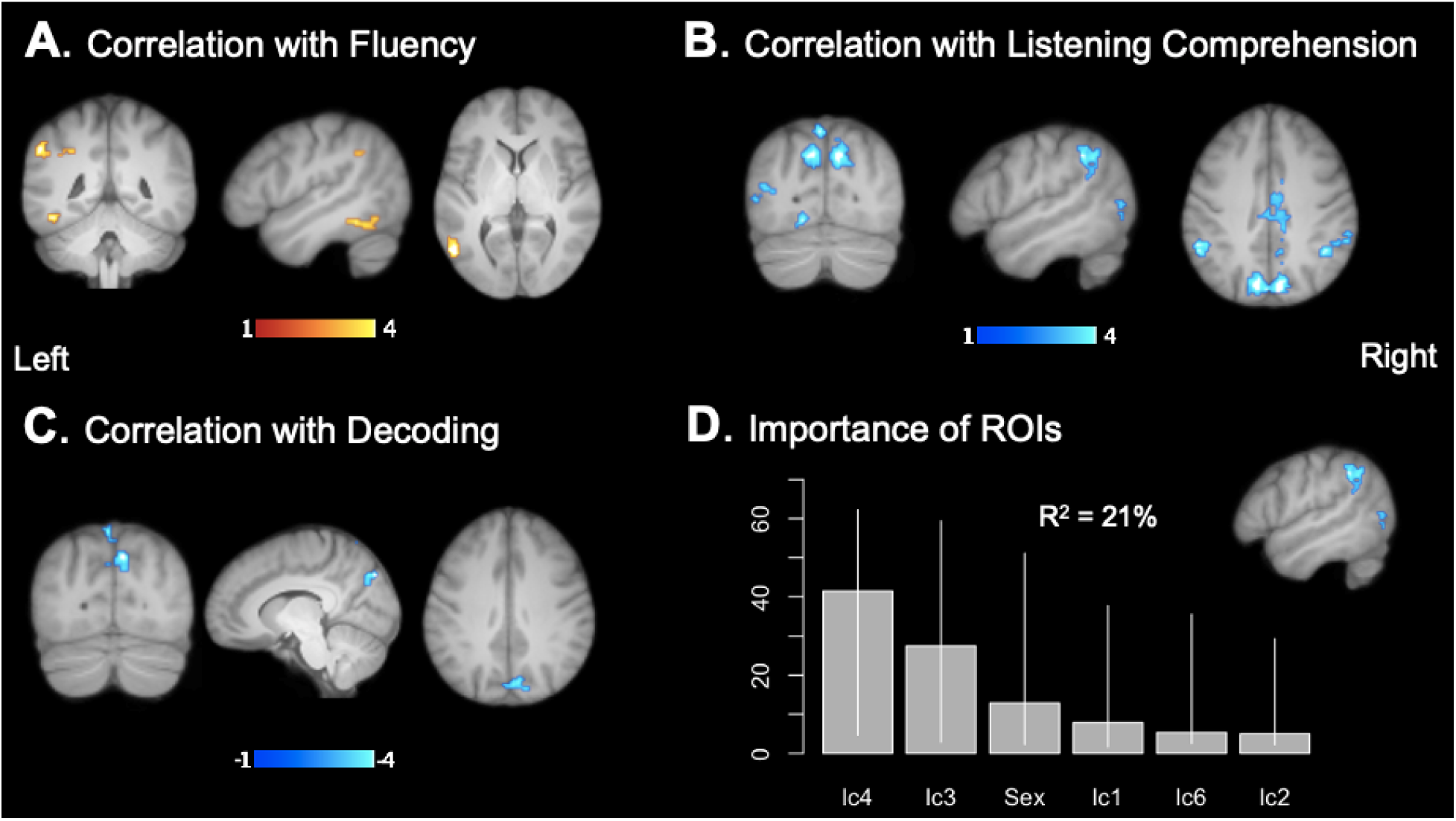
A-C: Regions in which there were significant correlations between activations (stories > arrows) and individual differences in performance on (A) Rapid Automatized Naming Letters (RANL); (B) Language Comprehension (LC), and (C) Word Attack (WA). Positive correlations in yellow/red and negative correlations in blue. Whole brain results for all participants. Z’s > 3.1, p’s < 0.05, corrected for multiple comparisons. (D) The two LC clusters (blue) that accounted for significant and unique variance in Reading Comprehension scores and a plot demonstrating the relative importance using the LMG method of each variable with 95% bootstrap confidence intervals.

The conjunction analysis for each pair of the constructs revealed overlapping regions of significant correlation between Language Comprehension and Decoding in posterior parietal regions. There were overlapping regions of correlated activation for Language Comprehension and Fluency in the left supramarginal/angular gyrus cluster and in a small middle temporal cluster. There were no significantly overlapping clusters between Fluency and Decoding.

**Table 2:**
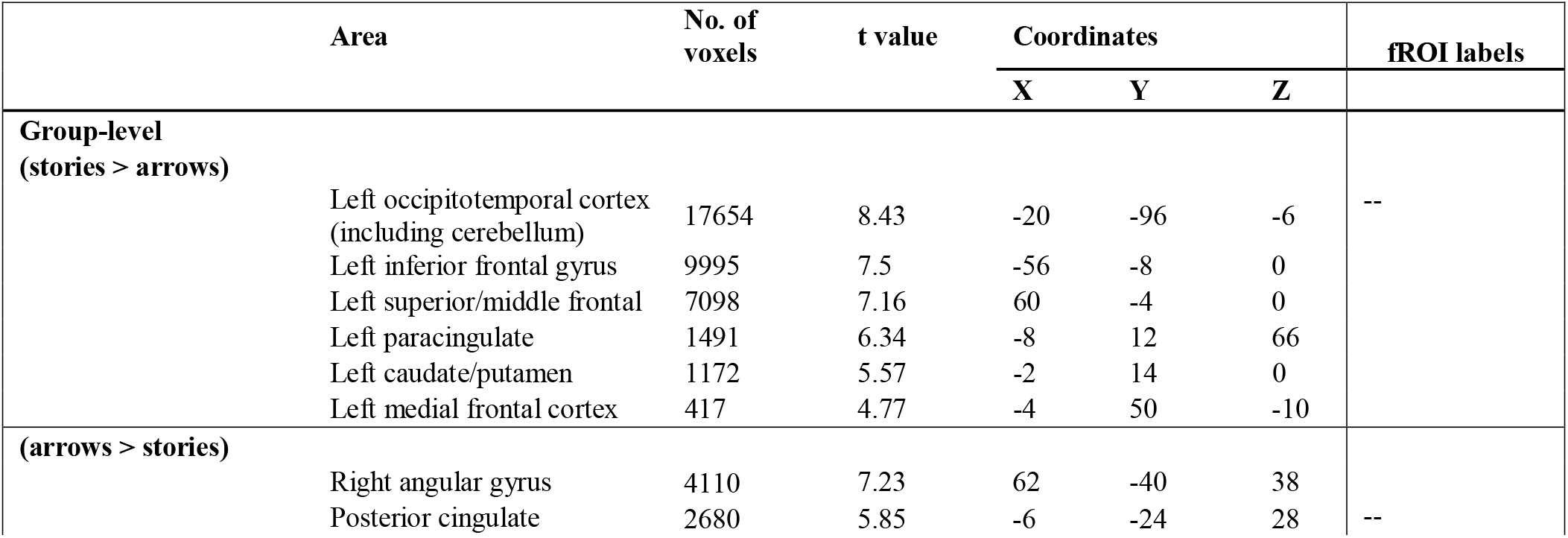
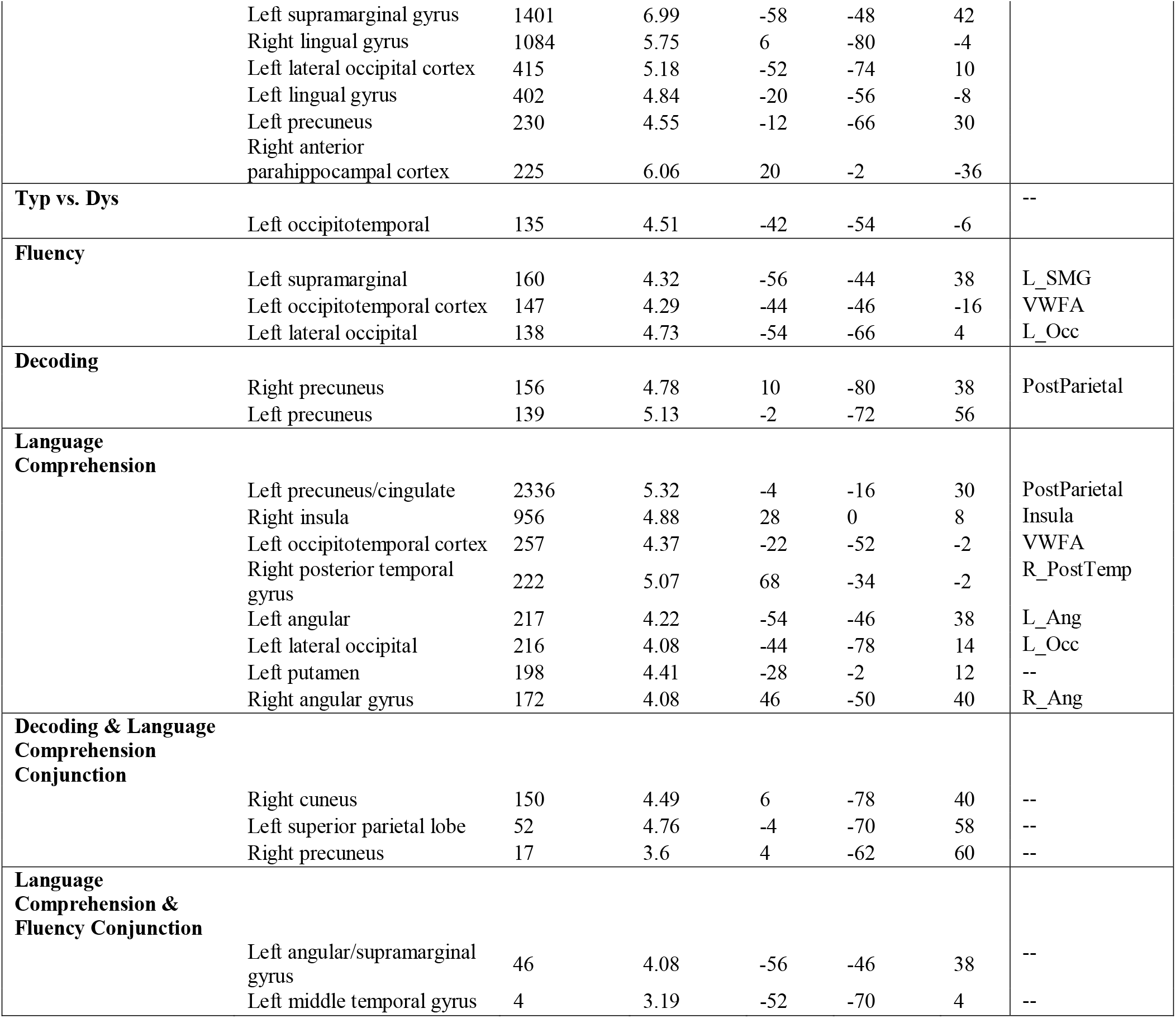
Regions of significant activation in stories > arrows whole-brain contrasts

| Area | No. of voxels | t value | Coordinates |  |  | fROI labels |
| --- | --- | --- | --- | --- | --- | --- |
|  |  |  | X | Y | Z |  |
| **Group-level (stories > arrows)** |  |  |  |  |  | -- |
| Left occipitotemporal cortex (including cerebellum) | 17654 | 8.43 | -20 | -96 | -6 |  |
| Left inferior frontal gyrus | 9995 | 7.5 | -56 | -8 | 0 |  |
| Left superior/middle frontal | 7098 | 7.16 | 60 | -4 | 0 |  |
| Left paracingulate | 1491 | 6.34 | -8 | 12 | 66 |  |
| Left caudate/putamen | 1172 | 5.57 | -2 | 14 | 0 |  |
| Left medial frontal cortex | 417 | 4.77 | -4 | 50 | -10 |  |
| **(arrows > stories)** |  |  |  |  |  | -- |
| Right angular gyrus | 4110 | 7.23 | 62 | -40 | 38 |  |
| Posterior cingulate | 2680 | 5.85 | -6 | -24 | 28 |  |

|  |  |  |  |  |  |  |  |
| --- | --- | --- | --- | --- | --- | --- | --- |
|  | Left supramarginal gyrus | 1401 | 6.99 | -58 | -48 | 42 |  |
|  | Right lingual gyrus | 1084 | 5.75 | 6 | -80 | -4 |  |
|  | Left lateral occipital cortex | 415 | 5.18 | -52 | -74 | 10 |  |
|  | Left lingual gyrus | 402 | 4.84 | -20 | -56 | -8 |  |
|  | Left precuneus | 230 | 4.55 | -12 | -66 | 30 |  |
|  | Right anterior parahippocampal cortex | 225 | 6.06 | 20 | -2 | -36 |  |
| **Typ vs. Dys** | Left occipitotemporal | 135 | 4.51 | -42 | -54 | -6 | -- |
| **Fluency** | Left supramarginal | 160 | 4.32 | -56 | -44 | 38 | L_SMG |
|  | Left occipitotemporal cortex | 147 | 4.29 | -44 | -46 | -16 | VWFA |
|  | Left lateral occipital | 138 | 4.73 | -54 | -66 | 4 | L_Occ |
| **Decoding** | Right precuneus | 156 | 4.78 | 10 | -80 | 38 | PostParietal |
|  | Left precuneus | 139 | 5.13 | -2 | -72 | 56 |  |
| **Language Comprehension** | Left precuneus/cingulate | 2336 | 5.32 | -4 | -16 | 30 | PostParietal |
|  | Right insula | 956 | 4.88 | 28 | 0 | 8 | Insula |
|  | Left occipitotemporal cortex | 257 | 4.37 | -22 | -52 | -2 | VWFA |
|  | Right posterior temporal gyrus | 222 | 5.07 | 68 | -34 | -2 | R_PostTemp |
|  | Left angular | 217 | 4.22 | -54 | -46 | 38 | L_Ang |
|  | Left lateral occipital | 216 | 4.08 | -44 | -78 | 14 | L_Occ |
|  | Left putamen | 198 | 4.41 | -28 | -2 | 12 | -- |
|  | Right angular gyrus | 172 | 4.08 | 46 | -50 | 40 | R_Ang |
| **Decoding & Language Comprehension Conjunction** | Right cuneus | 150 | 4.49 | 6 | -78 | 40 | -- |
|  | Left superior parietal lobe | 52 | 4.76 | -4 | -70 | 58 | -- |
|  | Right precuneus | 17 | 3.6 | 4 | -62 | 60 | -- |
| **Language Comprehension & Fluency Conjunction** | Left angular/supramarginal gyrus | 46 | 4.08 | -56 | -46 | 38 | -- |
|  | Left middle temporal gyrus | 4 | 3.19 | -52 | -70 | 4 | -- |

### Whole-Brain ROIs

Overall, 11 ROIs were extracted for the significant clusters identified in the whole-brain regression analyses (3 clusters for Fluency, 2 clusters for Decoding, and 6 clusters for Language Comprehension). All clusters were added into a regression model with Reading Comprehension as the predicted variable and Sex and Age as the control variables (Figure 3-D). For all variables, the variance inflation factor (VIF) values were below 10 except one of three Fluency clusters. This cluster, which was in the left supramarginal gyrus, was excluded from the regression models. The Shapiro–Wilk test revealed that some of the ROI values were not normally distributed (W’s < 0.95, *p*’s < .02). Accordingly, the lmPerm package (Wheeler, Torchiano, & Torchiano, 2016) in R was used to calculate permuted linear regression for all analyses. The R^2^ for the entire model was 0.21 (*p* < .001) and the best set of predictors included two of the Language Comprehension ROIs (lc3-left lateral occipital gyrus: 21.25% of R^2^; lc4-left angular gyrus: 42.74% of R^2^).

### Literature-based fROIs by Group

The Shapiro–Wilk test revealed that several of the fROI’s were not normally distributed and permuted linear regressions were calculated for all analyses with a stepwise approach to select the best model fit. VIF was evaluated for each model and clusters that exceeded the value of 10 were excluded from the particular model. For the Typ group, Fluency was significantly predicted by VWFA activation only, accounting for 10.37% of the variance (ηp^2^ = 0.12), with higher Fluency associated with increased activation in VWFA. Three fROIs had to be excluded in the Language Comprehension model due to high VIF: Insula, L_lOcc, and R_IFG. Language Comprehension was significantly predicted by Age (r^2^ = 0.13, ηp^2^ = .11), Sex (r^2^ = 0.28, ηp^2^ = 0.19), PostParietal (r^2^ = 0.28, ηp^2^ = 0.05), VWFA (r^2^ = 0.11, ηp^2^ = .06), R_Ang (r^2^ = 0.12, ηp^2^ = 0.19), and R_IFGorb (r^2^ = 0.04, ηp^2^ = 0.08). Increased recruitment of these regions was associated with lower Language Comprehension scores. For the Decoding model, R_IFGorb had to be excluded due to a high VIF. Decoding was significantly predicted by Age only (r^2^ = 0.17, ηp^2^ = 0.18).

For the Dys group, Fluency was significantly predicted by Age (r^2^ = 0.07, ηp^2^ = 0.004), Sex (r^2^ = 0.19, ηp^2^ = 0.25), and L_SMG (r^2^ = 0.1, ηp^2^ = 0.1). Better Fluency was associated with increased activation of this region. Several fROI’s had to be removed from the Language Comprehension model due to high VIF: PostParietal, Insula, R_SMG, and L_IFG. Language Comprehension was significantly predicted by L_IFG (r^2^ = 0.1, ηp^2^ = 0.04) and L_IFGorb (r^2^ = 0.18, ηp^2^ = 0.2). Better Language Comprehension was associated with increased activation of L_IFGorb and decreased activation of L_IFG. Due to high VIF, R_IFGob was removed from the Decoding model. Decoding was predicted by PostParietal (r^2^ = 0.09, ηp^2^ = 0.05), L_IFG (r^2^ = 0.04, ηp^2^ = 0.01), L_IFGob (r^2^ = 0.4, ηp^2^ = 0.48), and R_IFG (r^2^ = 0.16, ηp^2^ = 0.14). Increased activation of these regions was associated with lower Decoding.

**Figure 4:**
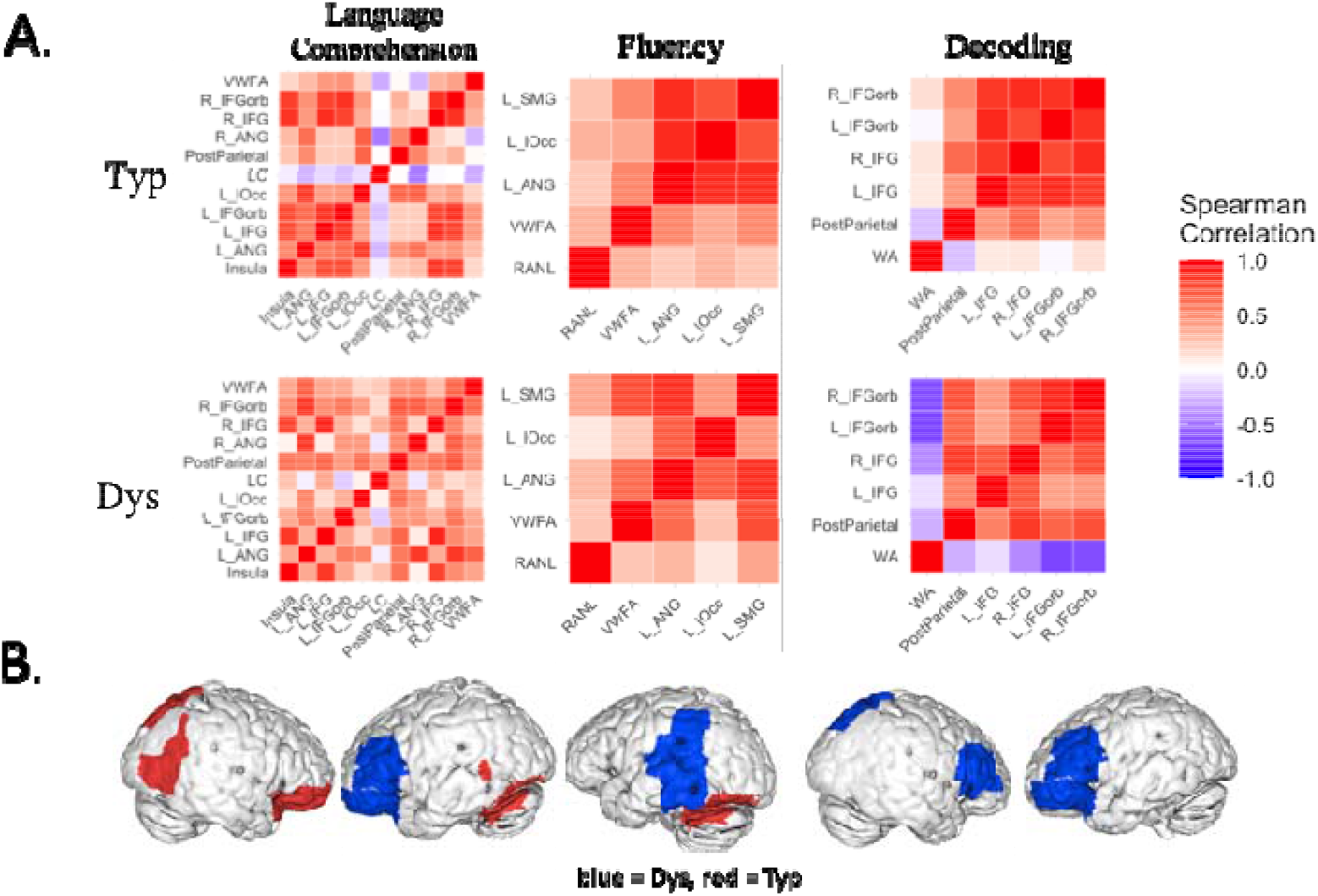
A) Correlations between Language Comprehension, Fluency, and Decoding scores and the fROIs and among the fROIs separately by group. B) fROIs that demonstrated a significant association in the stepwise regression model for each of the subskills for the Typ group (red), Dys group (blue).

To test which fROIs explained significant variance in reading comprehension performance in Dys and Typ groups, we included clusters that were significant in the six Decoding, Language Comprehension, and Fluency models in a stepwise regression model predicting Reading Comprehension separately in each group. In the Typ group, after removing several clusters due to high VIF (Insula, R_SMG, R_IFG), the overall model explained 49.58% of variance in Reading Comprehension scores with Sex explaining 30% of total variance (ηp^2^ = 0.03), with higher scores for males. The significant fROIs included: PostParietal explaining 14% (ηp^2^ = 0.21), L_IFGorb explaining 6% (ηp^2^ = 0.001), and L_IFG explaining 14% (ηp^2^ = 0.14). For Dys, after the high VIF variables were removed (L_SMG, Insula, R_SMG), the model accounted for 55.83% of total variance with the only significant predictor PostParietal explaining 25% of total variance (ηp^2^ = 0.13).

**Figure 5:**
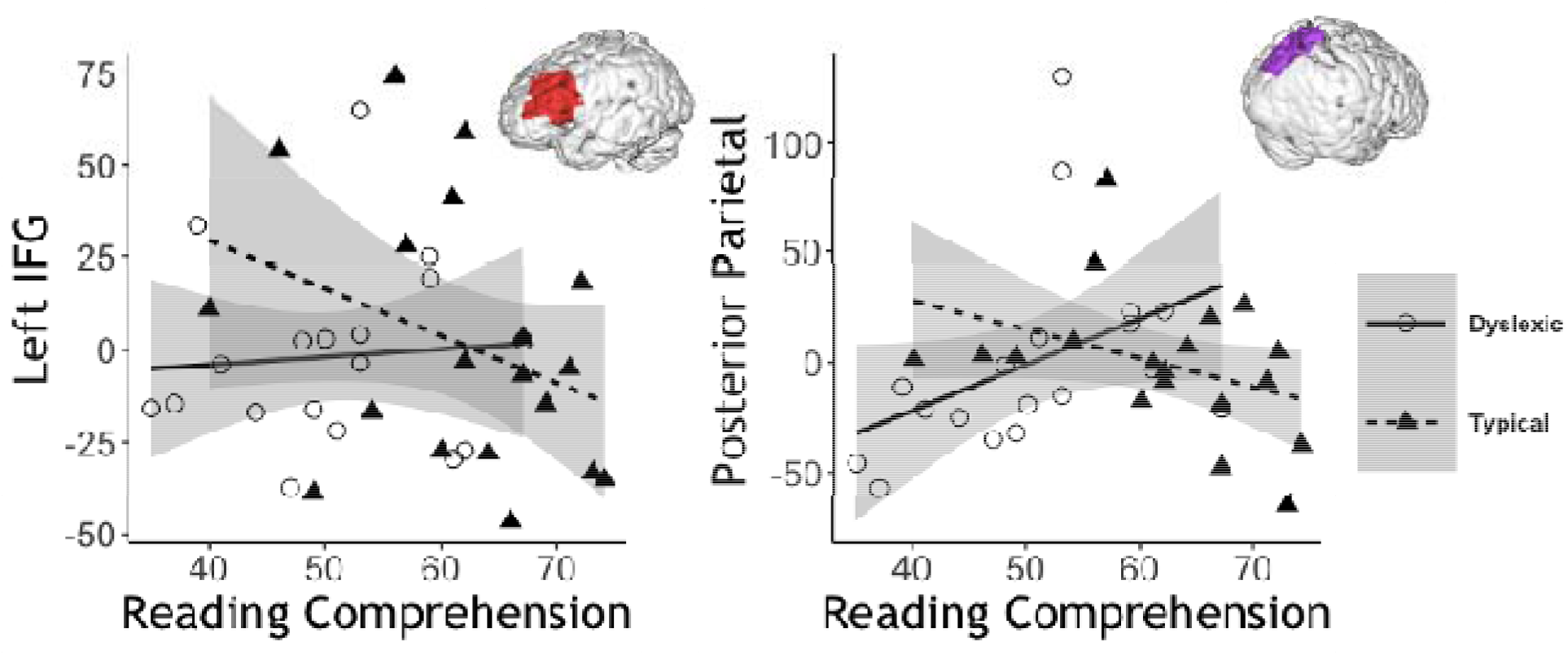
Association with reading comprehension for the best set of fROI predictors for Dys -- Posterior Parietal (right panel) and for Typ -- Posterior Parietal, Left IFG (left panel), and Left IFGorb (not shown).

## Discussion

The current study investigated, in adults with and without dyslexia, how brain activation during a naturalistic passage reading task was related to out-of-scanner components of reading comprehension as proposed by the Simple View of Reading: decoding accuracy, decoding fluency, and language comprehension. Consistent with the Simple View framework, the three constructs explained unique and significant variance in reading comprehension. We found distinctive patterns of activation in relation to individual differences in fluency and language comprehension. The activation related to decoding accuracy, however, was entirely overlapping with some of the regions associated with language comprehension. Activation in the inferior parietal and middle temporal language-comprehension ROIs made the largest contribution to reading comprehension. Importantly, confirming the validity of our fMRI paradigm, the stories > arrows contrast involved the expected dorsal and ventral reading regions. When examining the neural correlates of the three constructs in each of the groups using literature-based regions of interest, we found evidence that the neural substrates of comprehension differ in adults with and without dyslexia and the relative behavioral contribution of the three components to reading comprehension was different as well. Our results suggest that during naturalistic reading, individuals with dyslexia recruit cognitive and linguistic systems to support word decoding, thereby diverting resources from, and potentially impeding, comprehension processes. These findings reveal the underlying mechanisms of reading comprehension deficits of adults with dyslexia.

### Behavioral findings

The behavioral results confirmed the basic tenants of the Simple View framework and demonstrated a unique and significant contribution of fluency, decoding, and language comprehension to reading comprehension. The model that included the three variables in addition to sex and age accounted for 50% of variance in reading comprehension performance. This percentage represents the lower end of the range of explained variance reported in previous studies of adults (46-70% of variance; Braze et al., 2016; Landi, 2010; Talwar et al., 2018). Behavioral studies investigating reading comprehension components have commonly used latent factors drawing on multiple indicators to represent each of the three constructs, rather than single assessments. It is therefore possible that such factors explain more variance in reading comprehension than was explained in the current study. The sample size and the relatively limited battery of assessments in the current study precludes applying a factor analysis approach to identifying components of the decoding, fluency, and language comprehension subskills. Nevertheless, our behavioral findings support the validity of examining the three constructs in relation to neural activation during reading in adults.

There were significant differences in reading comprehension between the typical readers and individuals with dyslexia and the proportional contributions of fluency, decoding, and language comprehension components differed between the two groups. Consistent with the prediction of the Simple View framework that at higher levels of decoding, reading comprehension will be primarily explained by language comprehension, the only significant contributor to reading comprehension skills in typical readers was language comprehension. Indeed, it has been previously demonstrated that in higher grades and in adulthood, factors subsumed by language comprehension such as inference making, vocabulary, and background knowledge make large contributions to reading comprehension (see Ahmed et al., 2016). In contrast, in adults with dyslexia all three factors made significant contributions to explaining reading comprehension, with language comprehension explaining the largest percentage of variance. It has been suggested that the contribution of decoding to reading comprehension in adults with dyslexia is due to the slow and impaired word reading that creates a bottleneck for comprehension of texts because of additional cognitive demands (Crain & Shankweiler, 1990).

### Associations and dissociations among neural correlates of the Simple View of Reading

#### Decoding Correlates

Across all participants, we found a negative correlation between decoding accuracy (Word Attack scores) and activation in bilateral posterior parietal and precuneus regions. Separate analyses by group using a literature-based posterior parietal multi-domain fROI, revealed that the negative association between posterior parietal regions and decoding was driven by the dyslexic group only. Additional correlates of decoding in the dyslexic group included bilateral IFG and right IFG orbitalis. Posterior parietal/precuneus regions have been implicated in attentional modulation of complex tasks (Cavanna & Trimble, 2006). The precuneus has been frequently invoked in studies of language comprehension (Binder et al., 2009; Price, 2012) and reading (Chyl et al., 2018; Rimrodt et al., 2009; Roe et al., 2018; Ryherd et al., 2018; Schulz et al., 2008, 2009; Shaywitz et al., 2002), and there is evidence to suggest it represents the extent to which attentional resources are allocated to these tasks (Kuperberg et al., 2003; Schulz et al., 2008, 2009). Consistent with the current findings, several studies have demonstrated negative associations between activation in the precuneus and out-of-scanner reading (Chyl et al., 2018; Rimrodt et al., 2009; Ryherd et al., 2018). Cumulatively, these findings suggest that adults with poorer decoding skills invoked the attentional regions during reading to a greater extent than adults with better decoding skills. Importantly, the current study is the first study employing a connected-text paradigm in relation to individual differences in the subcomponents of reading. Previous studies that have identified the neural systems underlying reading in relation to decoding skills used single-word or sentence-reading tasks. The current findings, therefore, provide initial evidence of a relation between activation in the precuneus and individual differences in decoding skill as applied to reading connected text.

The increased importance of prefrontal regions for reading in individuals with dyslexia has been highlighted by previous studies (see Hancock et al., 2017). Several meta-analyses have documented increased activation in these regions in individuals with dyslexia relative to controls during phonological tasks (Maisog et al., 2008; Richlan et al., 2013). Hyperactivation in these regions has been interpreted as compensatory. In order to establish a compensatory role for these systems, however, it is important to demonstrate their engagement in relation to behavioral performance (Hancock et al., 2017). The few studies that have done this have demonstrated both positive and negative associations between bilateral inferior frontal regions and reading-related skills (Bach et al., 2010; Horowitz-Kraus et al., 2013; Ingvar et al., 2002; Patael et al., 2018; Rumsey et al., 1994; Ryherd et al., 2018). We found that individuals with worse Word Attack scores in the dyslexic group recruited the bilateral inferior frontal regions to a greater extent. The inferior frontal gyrus, bilaterally, represents the anterior language system involved in language comprehension and production, but has also been associated with domain-general cognitive and perceptual functions including attention, working memory, inhibitory control, planning/goal-directed behaviors, fluid intelligence, and consciousness (Hancock et al., 2017). Inferior frontal regions have been implicated in the resolution of phonological ambiguity via top-down modulation of activity in posterior phonological systems (Burton et al., 2000; Gow et al., 2008; Myers, 2007; Zatorre et al., 1996). These regions were also associated with better Language Comprehension skills in the dyslexic group. Together, the current findings provide novel evidence for the increased involvement of language comprehension and cognitive systems during reading to compensate for poor decoding skills in dyslexia.

#### Language Comprehension Correlates

We found increased activation of regions that are part of the cingulo-opercular networks (i.e. cuneus, cingulate, insula) as well as basal ganglia regions to be associated with worse language comprehension performance in the whole-brain analysis. These regions have been shown to support executive functions (e.g., inhibitory control, attentional selection, conflict resolution, maintenance and manipulation of task sets) for both linguistic and non-linguistic tasks (e.g., Duncan & Owen, 2000; Fedorenko et al., 2013; Hugdahl et al., 2015; see Fedorenko, 2014). The few previous studies investigating the neural correlates of individual differences in language comprehension also found increased engagement of these attentional and cognitive control systems during reading (Roe et al., 2018; Ryherd et al., 2018). Furthermore, cingulate and precuneus regions have been associated with updates in and integration of the reader’s mental model during discourse comprehension (Gernsbacher & Kaschak, 2003; Robertson et al., 2000; St George et al., 1999; Whitney et al., 2009; Yarkoni, Speer, & Zacks, 2008a).

Additionally, there was a negative correlation with clusters of activation in the bilateral middle temporal gyrus and angular gyrus. These regions are thought to be part of a distributed semantic system across modalities (e.g., speech and text) and stimuli (language, faces) (Hartwigsen et al., 2015; Price, 2012; Price et al., 1997). The engagement of the semantic network for reading as a function of comprehension skills is in accordance with the behavioral literature linking poor comprehension to deficits in semantic processing (see Landi & Ryherd, 2017). The current findings, however, indicate a negative association between comprehension skills and the engagement of this network, showing increased recruitment of these regions in participants who had poorer language comprehension. These findings are consistent with several previous studies that found a negative association between word reading skill and the engagement of the semantic network (Rimrodt et al., 2009; Welcome & Joanisse, 2012), but not with other studies that found a positive association (Aboud et al., 2016; Ettinger-Veenstra et al., 2016; Ryherd et al., 2018). For example, one study found decreased activation for skilled comprehenders in regions including angular gyrus and middle temporal gyrus during word reading (Welcome & Joanisse, 2012). In another study, there was a positive association between comprehension skills and activation in middle temporal regions and a negative association in inferior parietal regions during sentence-by-sentence passage reading (Ryherd et al., 2018). Therefore, our findings extend the previous literature in demonstrating that during naturalistic reading adults with lower comprehension skills engage both the domain-general executive systems and the language-specific semantic system to a greater extent.

When examining the Language Comprehension correlates by group, however, the semantic system identified in the whole-brain analysis was found to be negatively associated with comprehension skills in the typically reading group only. Furthermore, the posterior parietal area was also recruited to a greater extent in the typical adults with poor comprehension skills, but not in the dyslexic adults. In the dyslexic group, better Language Comprehension skills were associated with increased activation of the left IFG only. Thus, the domain-general posterior cognitive system and the semantic system were deployed to support reading in adults with poor comprehension but adequate decoding.

#### Fluency Correlates

Across all participants, better Fluency was positively associated with increased activation of the left ventral occipital-temporal region and left temporoparietal regions. In the typically reading group, better Fluency was associated with increased activation of the putative visual word form area (VWFA), a region that develops specialization for automatic word recognition with increased reading experience (Centanni et al., 2018; Kronbichler et al., 2004; McCandliss et al., 2003). Accordingly, previous studies have demonstrated an increased recruitment of VWFA with increased reading fluency demands (Benjamin & Gaab, 2012; Langer et al., 2015). In the Dys group, better Fluency skills were associated with increased recruitment of the left temporoparietal systems, comprising the dorsal phonological reading network as well as the semantic systems. The posterior middle temporal region has been implicated in previous studies as an additional region of difference in fluency performance (Meyler et al., 2007; Roe et al., 2018).

The differentiation between the two groups in the neural correlates of fluency is consistent with developmental theories of neural specialization for reading (Pugh et al., 2000; Sandak et al., 2004; Younger et al., 2017). Less skilled readers engage the dorsal temporoparietal network for phonological analysis and recoding, whereas skilled readers rely on the rapid, ventral stream for orthographic computations. The automaticity of the ventral system facilitates reading fluency which in turn is thought to lead to better reading comprehension. This theory is supported by metanalytic findings showing a more consistent recruitment of the ventral system during reading in adults as compared to children (Martin et al., 2015). Together, the current results support the notion that RAN, as a measure of fluency, represents the automaticity of integration across the individual components of the reading circuit: the orthographic, phonological, and semantic systems (Norton & Wolf, 2011). In individuals with dyslexia, fluency depends on more efficient utilization of the dorsal system, but in typical readers, increased fluency depends on the automaticity of the ventral system.

#### Decoding & Language Comprehension

We examined whether there were commonly activated voxels for decoding and language comprehension skills. We found that the regions associated with decoding completely overlapped with the regions associated with language comprehension. Specifically, poor decoding and poor comprehension across all participants were associated with increased engagement of the domain-general posterior cognitive system regions previously demonstrated to be involved in cognitive control during reading (Horowitz-Kraus et al., 2013; Roe et al., 2018). Interestingly, in the dyslexic group the posterior parietal cognitive-control fROI was more activated in reading with decreasing decoding skills, but in the typically reading group the region was more involved with decreased language comprehension skills. The overlap in neural correlates of Language Comprehension and Decoding were also evident in left inferior frontal regions, which supported both decoding and comprehension in the Dys group. These results extend behavioral findings in support of the idea that both cognitive and linguistic resources are deployed at different stages during reading based on dyslexia status: in individuals with dyslexia they support lower-level word identification and in typical readers they support higher-level comprehension.

### Semantic regions explaining reading comprehension

We found that even though behaviorally the three constructs made significant and unique contributions to reading comprehension, only two semantic regions, left angular gyrus and middle temporal gyrus, explained a significant variance in these skills. Deficits in the semantic domain have been most consistently implicated in poor oral comprehension both in neuroimaging and in behavioral literature (see Landi & Ryherd, 2017). Furthermore, the developmental behavioral literature suggests that the unique contribution of decoding skills to variance in reading comprehension decreases across development and with increased mastery of reading (Catts et al., 2005; Hoover & Gough, 1990; Schatschneider et al., 2004). Instead, in these studies, the greatest contribution to reading comprehension was made by language comprehension skills as well as the shared variance between decoding and comprehension. Our neuroimaging findings in adults, therefore, mirror these behavioral results with the caveat that due to our enrollment criteria for dyslexia, the decoding skills in the current sample are oversampled on the lower end of the skill distribution. Therefore, the significant behavioral contribution of decoding to reading comprehension in our models could be due to the severity of the decoding deficits in our sample. Nevertheless, this is the first study to directly compare the relative contribution of the different neural components of reading to out-of-scanner reading comprehension. The findings provide the first neural evidence for the crucial role of the semantic network, above phonological and orthographic systems, for reading comprehension in adults.

### Differences in neural correlates of SVR in dyslexia

An important implication of the Simple View theory is that the interactive relationship between language comprehension and word reading influences reading comprehension. This interactivity implies that the relative contribution of the individual components to reading comprehension varies based on the levels of each of the component skills. Based on this formulation, in individuals with dyslexia, a greater relative contribution of decoding to reading comprehension than in typical readers would be expected. Our behavioral findings support this prediction. Furthermore, our neural findings provide important insights into the underlying mechanisms of the Simple View of Reading in dyslexia. In typical readers, consistent with the behavioral results, there were no significant differences in activation patterns in relation to decoding skills. In adults with dyslexia, both linguistic and cognitive systems were deployed to support decoding. The inferior frontal language system was recruited in individuals with lower decoding and higher comprehension skills, suggestive of increased linguistic top-down modulation of word identification in dyslexia, in order to compensate for the impairments in the posterior left-hemispheric reading regions.

In addition to the significant contribution of the semantic system to reading comprehension across all participants, the patterns of neural contributions to explaining variance in reading comprehension skills differed between the two groups. In the typically reading group, increased recruitment of the posterior parietal and left inferior frontal regions was associated with worse reading comprehension skills. In the dyslexic group, the multi-domain posterior parietal region made the only significant contribution to explaining differences in reading comprehension. Increased recruitment of this system was associated with better reading comprehension, but it was also recruited in relation to poor decoding skills in the dyslexic group. This suggests that the contribution of the cognitive mechanisms to explaining reading comprehension in adults with dyslexia is modulated through the recruitment of this system for decoding.

Reading comprehension occurs when lower-level processes, such as word decoding, are integrated with higher-level comprehension systems (Perfetti et al., 2008; Perfetti & Roth, 1980) These processes take place within a domain-general cognitive system with limited processing resources. Strong decoding skills free up cognitive capacity for higher-level comprehension of text and for processes related to formulating inferences while reading (Thurlow & Broek, 1997). Our findings offer insight into neural mechanisms of poor reading comprehension in adults with dyslexia, possibly indicating that this weakness could be due to insufficient cognitive resources to support linguistic comprehension when these resources are instead recruited to support accurate reading. The crucial role of these domain-general cognitive processes for comprehension is buttressed by extensive behavioral evidence of cognitive impairments in individuals with poor comprehension across multiple domains (e.g., planning, working memory, response inhibition, task switching) (Cutting et al., 2009; Locascio et al., 2010; Potocki et al., 2017; Protopapas et al., 2007; Sesma et al., 2009). Furthermore, evidence from poor readers supports the specificity of cognitive impairments for tasks involving reading, but not for other tasks (Roe et al., 2018). Our neuroimaging results therefore lend support to the proposal that in poor readers under naturalistic reading conditions, cognitive control resources are strained by word decoding, creating a bottleneck and weakening the contribution of these resources towards integration and comprehension (Hudson et al., 2005; Pikulski & Chard, 2005).

**Figure 6:**
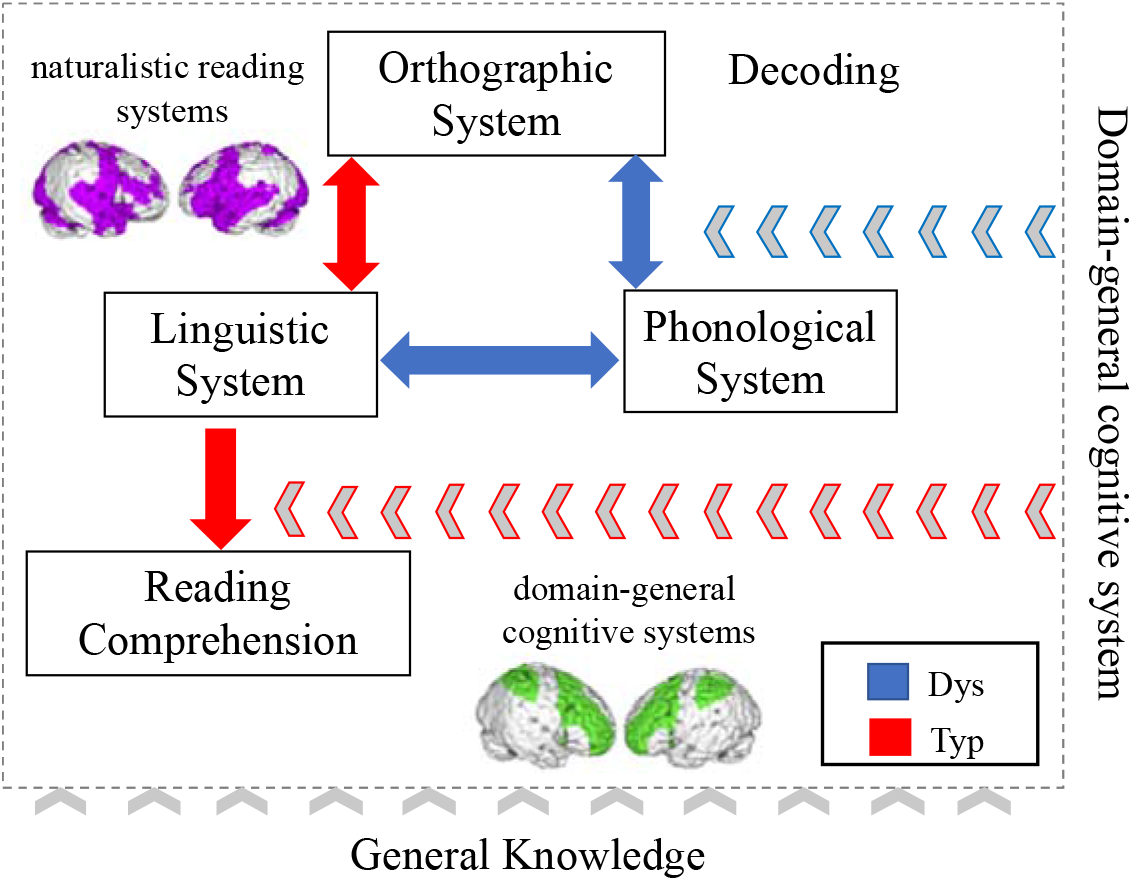
A diagram adapted from Perfetti & Stafura, 2014 shows the simplified framework of the process of text comprehension for typical readers (in red) and for individuals with dyslexia (blue), supported by the current neuroimaging evidence. In contrast to the original model, Linguistic System in the current model refers to the non-phonological linguistic processes (e.g., semantic, syntactic). The figure represents the process of reading from word processing to reading comprehension. In typical readers word identification is automatic and once word meaning is retrieved, cognitive resources (represented as the domain-general cognitive networks) are recruited to support linguistic comprehension. Alternatively, in individuals with dyslexia, word identification is not automatic and cognitive resources are deployed to support processes related to decoding (mapping words to their phonemic representations), subsequently compromising processes related to comprehension.

### Conclusions and Implications

Our findings provide a snapshot of the neural underpinnings of naturalistic reading and show how these mechanisms are related to individual abilities across different reading comprehension components based on the Simple View of Reading. It is possible that the central role of domain-general networks during reading in the current study is the result of the expository nature of the text used for the fMRI task. There has been some evidence showing that expository texts are more difficult to process (Best et al., 2004; Graesser & McNamara, 2011; McNamara et al., 2003) and place higher demands on executive functions networks than narrative texts, particularly in terms of inferencing and planning/organizing information (Baretta et al., 2009; Eason et al., 2012; Miller et al., 2013). The Simple View of Reading does not account for the interaction between reader differences and text characteristics and therefore these were not examined in the current study. Future studies are needed to delineate neural systems supporting expository versus narrative text types in relation to individual reader skills.

Additionally, as discussed above, the nature of the interaction among the three components changes across development, with greater independence of decoding and language comprehension found in older readers. The contributions of vocabulary and background knowledge to comprehension also increase with age and text difficulty. Furthermore, although there is convincing evidence for the robustness of the Simple View model across orthographies, the relative influence of decoding and language comprehension on reading comprehension varies across orthographies (Florit & Cain, 2011). In more transparent orthographies, fluency plays a greater role than in English. This suggests that our findings are limited to inference about English-speaking adults and that developmental studies across orthographies are needed to examine whether similar patterns of neural correlations are evident in children. Further, future studies may benefit from larger samples that support additional analyses and that may reveal more about variance within and between typically developing and dyslexic readers.

Despite the need for further investigations, the current study is the first to identify the neural correlates of the Simple View framework, as they unfold during naturalistic and typical reading. The Simple View has been a predominant framework of reading comprehension, and the current findings support the application of the model to dyslexia and reveal different behavioral and neural contributions to reading comprehension in this disorder.

## Data and Code Availability

The behavioral and fROI data and the analysis code used to support the findings of this study have been deposited in a Github repository (https://github.com/oozernov/neural_correlates_svr)

## Notes

We thank our participants. We thank the Athinoula A. Martinos Imaging Center at the McGovern Institute for Brain Research (MIT) and Atshusi Takahashi and Steve Shannon for data collection technical support. We thank Marina G. Monsivais, Sehyr Khan, and Karolina Wade for scoring the behavioral data.

## Supplemental Materials

**Supplemental Table 1:**
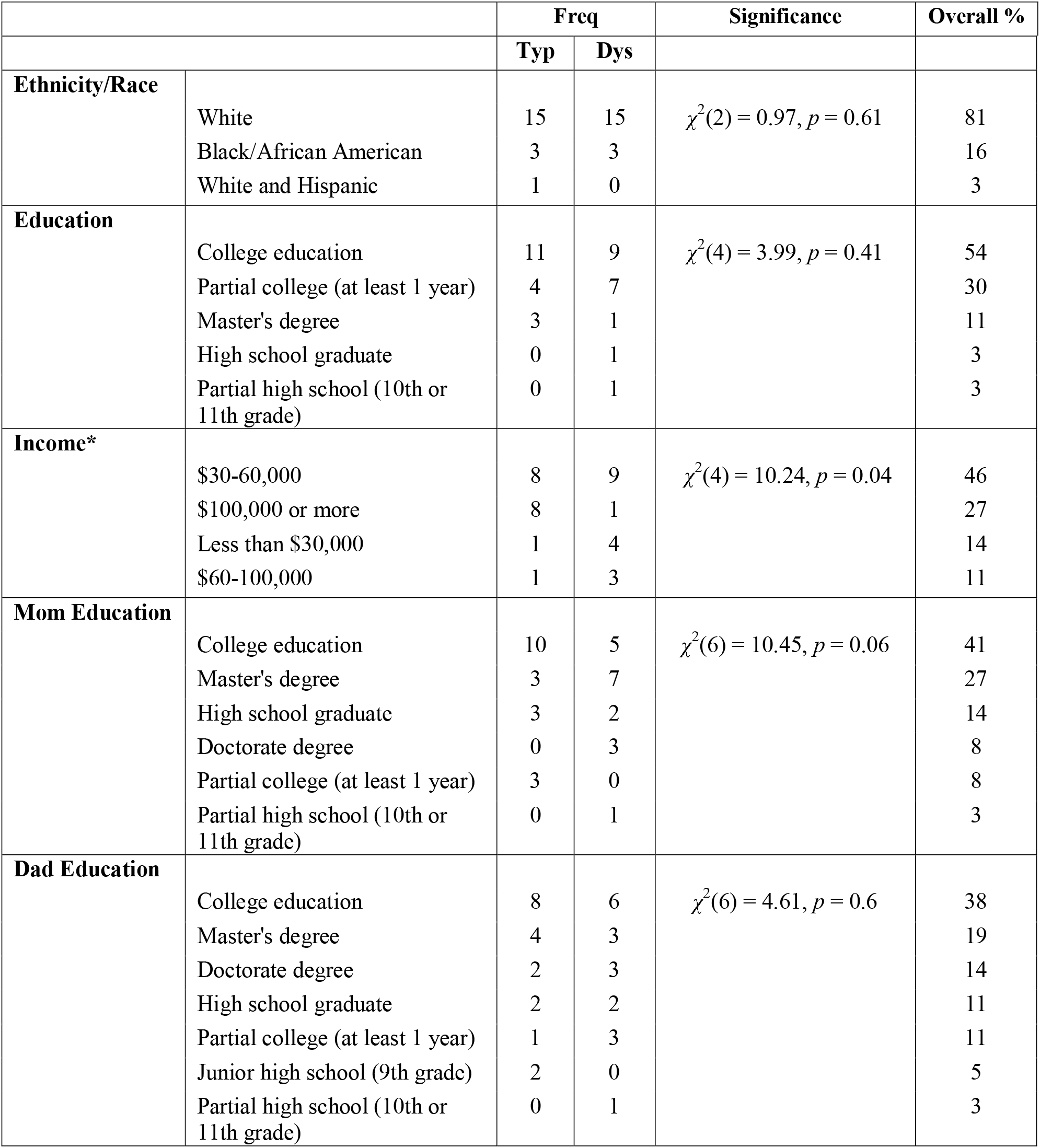
Participant demographic information

|  |  | Freq |  | Significance | Overall % |
| --- | --- | --- | --- | --- | --- |
|  |  | Typ | Dys |  |  |
| **Ethnicity/Race** | White | 15 | 15 | $\chi^2(2) = 0.97, p = 0.61$ | 81 |
|  | Black/African American | 3 | 3 |  | 16 |
|  | White and Hispanic | 1 | 0 |  | 3 |
| **Education** | College education | 11 | 9 | $\chi^2(4) = 3.99, p = 0.41$ | 54 |
|  | Partial college (at least 1 year) | 4 | 7 |  | 30 |
|  | Master's degree | 3 | 1 |  | 11 |
|  | High school graduate | 0 | 1 |  | 3 |
|  | Partial high school (10th or 11th grade) | 0 | 1 |  | 3 |
| **Income\*** | \$30-60,000 | 8 | 9 | $\chi^2(4) = 10.24, p = 0.04$ | 46 |
| | \$100,000 or more | 8 | 1 | | 27 |
| | Less than \$30,000 | 1 | 4 | | 14 |
| | \$60-100,000 | 1 | 3 | | 11 |
| **Mom Education** | College education | 10 | 5 | $\chi^2(6) = 10.45, p = 0.06$ | 41 |
|  | Master's degree | 3 | 7 |  | 27 |
|  | High school graduate | 3 | 2 |  | 14 |
|  | Doctorate degree | 0 | 3 |  | 8 |
|  | Partial college (at least 1 year) | 3 | 0 |  | 8 |
|  | Partial high school (10th or 11th grade) | 0 | 1 |  | 3 |
| **Dad Education** | College education | 8 | 6 | $\chi^2(6) = 4.61, p = 0.6$ | 38 |
|  | Master's degree | 4 | 3 |  | 19 |
|  | Doctorate degree | 2 | 3 |  | 14 |
|  | High school graduate | 2 | 2 |  | 11 |
|  | Partial college (at least 1 year) | 1 | 3 |  | 11 |
|  | Junior high school (9th grade) | 2 | 0 |  | 5 |
|  | Partial high school (10th or 11th grade) | 0 | 1 |  | 3 |

**Supplemental figure 1:**
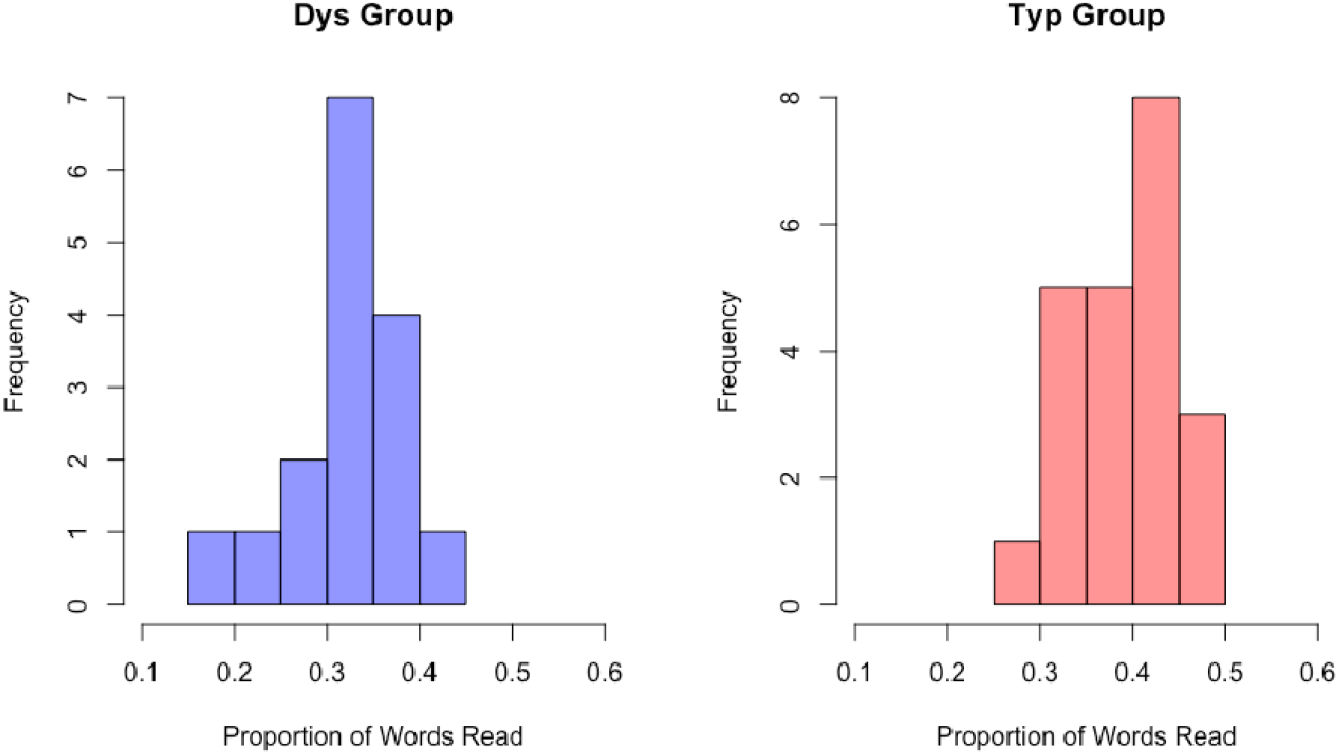
Proportion of words read during the in-scanner task by group

**Supplemental figure 2:**
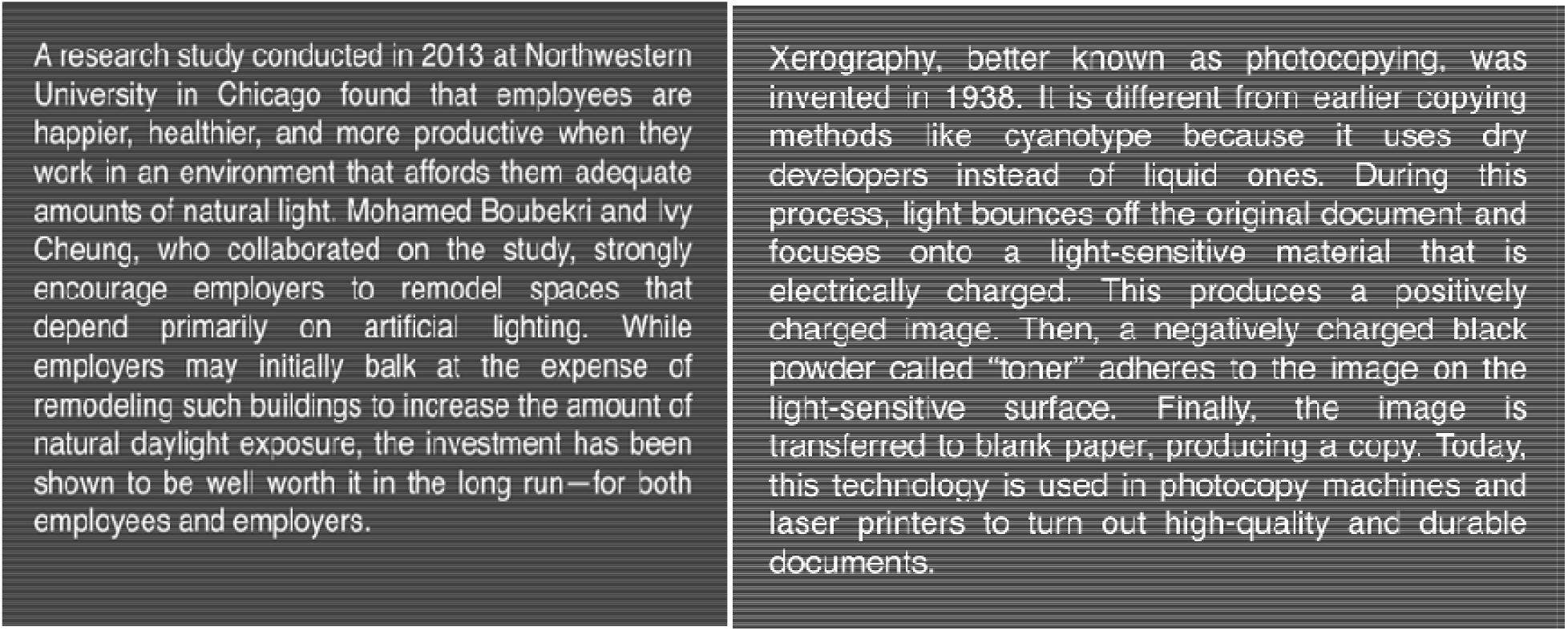
Example stories from the in-scanner reading task.

